# BiomarkerML: A cloud-based proteomics ML workflow for biomarker discovery

**DOI:** 10.1101/2025.10.16.682839

**Authors:** Yuhan Zhou, Anand K. Maurya, Yun Deng, Micah P. Fletcher, Connia Ren, Avigail Taylor

**Affiliations:** Institute of Molecular and Computational Medicine, Nuffield Department of Medicine, University of Oxford, Old Road Campus, Oxford, OX3 7BN

**Keywords:** machine learning, cloud-based workflow, classification and regression, proteomic biomarker identification

## Abstract

**Background:** High-throughput affinity and mass-spectrometry-based proteomic studies of large clinical cohorts generate high-dimensional proteomic data useful for accelerated disease biomarker discovery. A powerful approach to realizing the potential of these big, complex, and non-linear data, whilst ensuring reproducible results, is to use automated machine learning (ML) and deep learning (DL) pipelines for their analysis. However, there remains a gap in comprehensive ML workflows tailored to proteomic biomarker discovery and designed for biomedical researchers who need pipelines to optimally self-configure and automatically avoid over-fitting.

**Findings:** We present BiomarkerML, a cloud-based workflow for automated, reproducible, and efficient ML/DL analysis of proteomic data for biomarker discovery, designed for novice-ML users and implemented in Python, R and Workflow Description Language (WDL). BiomarkerML: ingests proteomic and clinical data alongside sample labels; pre-processes data for model fitting and optionally performs dimensionality reduction and visualization; fits a catalogue of ML and DL classification and regression models; and calculates model performance metrics for model comparison. Next, the workflow applies mean SHapley Additive exPlanations (SHAP) to quantify the contribution of each protein to model predictions across all samples. Finally, proteins with high mean SHAP values, and their co-expressed protein network interactors, are identified as candidate biomarkers. Importantly, hyperparameters - configuration variables set prior to training models - are automatically fine-tuned via grid-search, and BiomarkerML employs weighted, nested cross-validation to avoid model over-fitting and data leakage.

**Conclusions:** BiomarkerML is scalable, provides a standardized, user-friendly interface, and streamlines analyses to ensure reproducibility of results. Overall, BiomarkerML is a significant advancement, enabling novice-ML researchers to use cutting-edge ML/DL tools to identify disease biomarkers in complex proteomic data.

## Introduction

Mass spectrometry (MS) and affinity-based proteomic technologies such as Olink and SomaScan [1,2] have accelerated the field of biomarker discovery by making it possible to simultaneously measure thousands of proteins across many samples from large clinical cohorts. This is particularly important as disease biomarkers play a critical role in early diagnosis, prognosis, and therapeutic intervention [3,4]. However, the size, complexity and non-linearity of proteomic data present substantial analytical and interpretive challenges [5,6]. A powerful approach to addressing these challenges and realizing the potential of these data, whilst ensuring reproducible results, is to use automated, computationally efficient, machine learning (ML) and deep learning (DL) workflows tailored to proteomic data analysis [7,8]. However, despite rapid progress in the performance of ML and DL algorithms for biomarker discovery in proteomic data [9], the challenge of orchestrating these complex computational pipelines remains a significant barrier to uptake by biomedical researchers. Developing a comprehensive ML workflow tailored to proteomic biomarker discovery that implements ML best practices through an accessible interface, is key to making these methods usable by biomedical researchers [9,10]. Here, we present BiomarkerML, a cloud-based workflow for automated, reproducible, and efficient ML/DL analysis of proteomic data for biomarker discovery, designed for novice-ML users and implemented in Python, R and Workflow Description Language (WDL).

To ensure that complex machine learning analyses are both accessible and rigorous, BiomarkerML integrates four high-level methodological facets essential for these goals, yet often underrepresented in existing pipelines. First, we implement a class-weighted nested cross-validation framework with inner and outer resampling loops, which mitigates selection bias, reduces data leakage, and provides robust model performance estimates, particularly critical for high-dimensional and class-imbalanced proteomic data [11,12]. Second, we automate hyperparameter optimization using scalable search algorithms within the nested design, removing the burden of manual tuning while ensuring reproducibility [13,14]. Third, we incorporate model-agnostic mean SHapley Additive exPlanations (SHAP) values, computed by averaging SHAP values across all samples, to quantify protein-level feature importance, thereby improving model interpretability and enabling transparent biomarker identification [15,16]. Finally, recognising that any list of putative biomarkers identified using purely ML/DL methods will always be the smallest set of explainers for the classification or regression problem posed, we use protein network analyses to expand the list of putative biomarkers to include proteins with low mean SHAP values, but which are biologically related to high-mean-SHAP-value proteins [17,18]. Together, these components address common pitfalls in proteomics machine learning workflows and provide a unified solution that supports both linear and non-linear modelling strategies across a range of ML and DL algorithms.

To realize these facets, BiomarkerML implements the following components:

1. **Data pre-processing**: BiomarkerML handles missing values and normalizes data across heterogeneous measurement platforms and modalities.
2. **Data visualization**: The workflow also provides interactive visualizations using dimensionality reduction techniques (e.g., PCA, UMAP, t-SNE) to explore data structure.
3. **Catalogue of ML and DL models**: BiomarkerML includes both established and state-of-the-art ML and DL models suitable for binary classification, multiclass classification, and regression, enabling its application to a range of biological and clinical questions.
4. **Class-weighted, nested cross-validation**: The workflow employs class-weighted nested cross-validation with inner and outer loops to ensure unbiased model evaluation [11,12,19].
5. **Dimensionality reduction and feature selection**: BiomarkerML provides linear and non-linear methods for dimensionality reduction and feature selection, which can be combined with models that are not well suited to high dimensionality proteomic data.
6. **Automated hyperparameter tuning**: Hyperparameters are model configuration variables that must be tuned prior to training models; BiomarkerML tunes these automatically.
7. **Performance metric visualization**: Our workflow compares models using standard classification and regression metrics [19].
8. **Mean SHAP-based biomarker identification**: BiomarkerML uses mean SHAP values to quantify the contribution of each protein to model predictions, enabling consistent comparison of feature importance across a wide range of ML and DL models [15,16].
9. **Protein network analysis for further biomarker detection:** The workflow also conducts protein co-expression and interaction network analyses of high-mean-SHAP-value proteins to identify first-degree biologically related proteins as an expanded list of putative biomarkers.
10. **Report generation:** Finally, BiomarkerML collates and summarises results in an automatically generated report. Results are also output as tables, in comma-separated-value format, for ingestion into downstream analytical pipelines.

Existing tools incorporate subsets of these components, but BiomarkerML is the first to integrate all ten into a unified solution for proteomic biomarker discovery, with some components rarely implemented elsewhere (see Supplementary Section 1, Table S1). Importantly, rather than aggregating pre-existing models or relying on black-box ML tools, as is the case, for example, in the MLme workflow [20], we have implemented all machine learning and deep learning models natively in Python using *scikit-learn* and *PyTorch*, thus maximizing transparency, reproducibility, and extensibility of the software. Notably, we present two custom-designed classification architectures based on Variational Autoencoders (VAEs)—a class of generative AI models within the broader deep learning paradigm—specifically optimized to address the unique challenges of proteomic data, including high dimensionality, non-linearity, and multicollinearity. [21,22]. Overall, BiomarkerML is a modular, fully automated, end-to-end workflow implemented in Python, R, and WDL. By integrating rigorous evaluation, automated hyperparameter optimization, model interpretability, and biological contextualization, the workflow enables researchers—regardless of ML expertise—to perform scalable, reproducible analyses of complex proteomic datasets for biomarker discovery and therapeutic target prioritization.

## Key Features of BiomarkerML

### Robust Multi-Stage Pre-processing and Advanced Dimensionality Reduction

BiomarkerML offers a carefully engineered, multi-stage data pre-processing pipeline that integrates standardized feature scaling and removal of missing data, followed by one of seven complementary dimensionality reduction techniques [23] to address the high dimensionality and non-linearity of proteomic data (see Fig. 1 for an overview of the workflow, and Supplementary Section 2 for methodological details). These are: Principal Component Analysis (PCA) [24], Uniform Manifold Approximation and Projection (UMAP) [25], t-distributed Stochastic Neighbour Embedding (t-SNE) [26], Kernel Principal Component Analysis (KPCA) [24], Partial Least Squares (PLS) regression and ElasticNet regularization [27]. These techniques are embedded within a nested cross-validation framework, ensuring that critical biological signals are retained while overfitting is minimized, thereby enhancing both the generalizability and interpretability of downstream predictive models [28].

**Figure 1:**
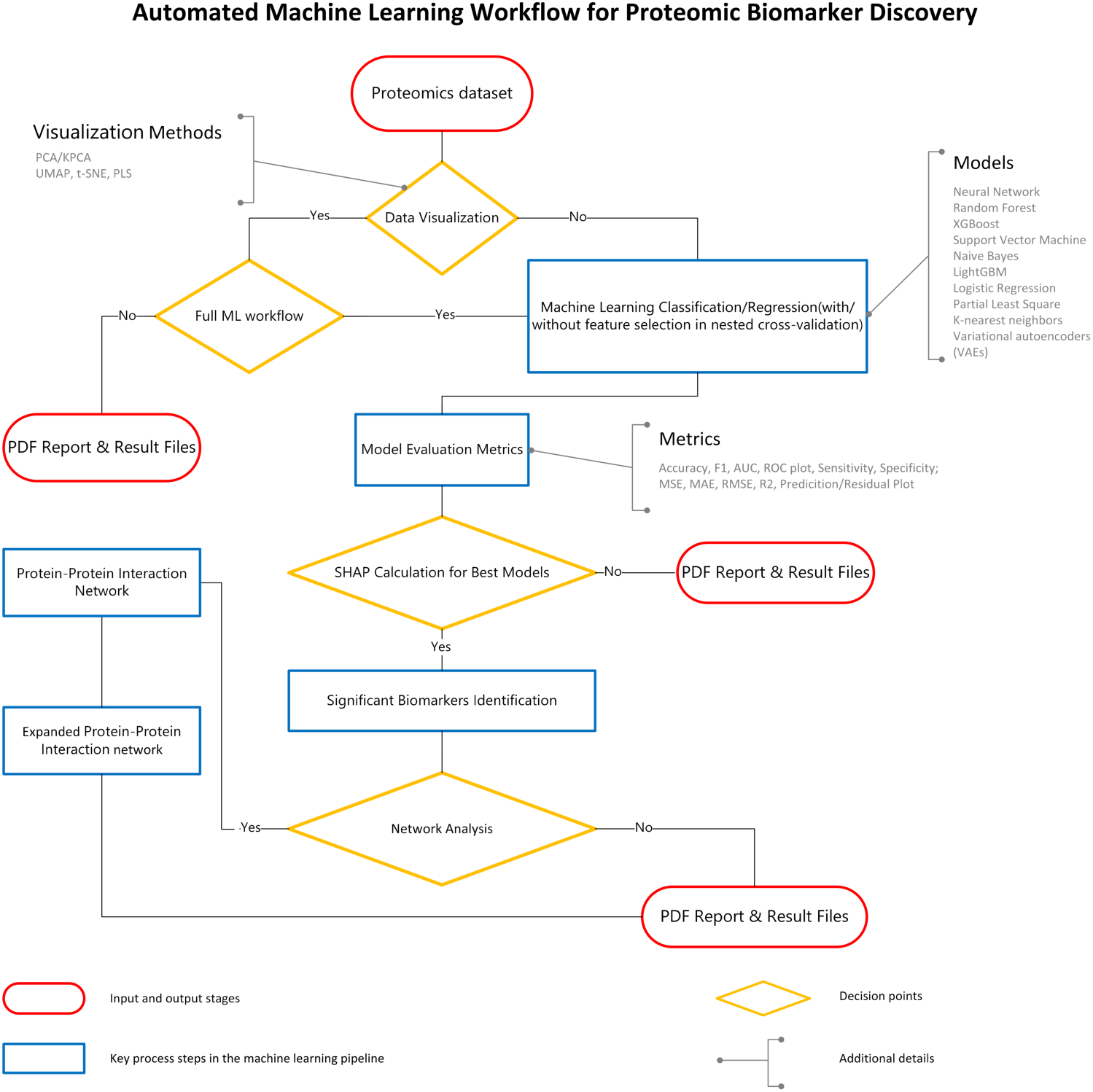
Overview of the BiomarkerML workflow.

### Comprehensive Model Comparison with Automated Hyperparameter Optimization

To navigate the inherent complexity and heterogeneity of proteomic datasets, BiomarkerML supports a broad range of classical machine learning algorithms such as Random Forest (RF), Support Vector Machines (SVM), Extreme Gradient Boosting (XGBoost), and LightGBM in *Scikit-learn*, as well as two original deep learning Variational Autoencoder (VAE) architectures custom-built in *PyTorch* [29,30]. Hyperparameter tuning is performed automatically using Bayesian optimization via *Optuna* within a nested cross-validation framework [13]. This enables a robust, direct comparison of model performance which is assessed using accuracy, F1-score, sensitivity, specificity, and Area Under the Curve - Receiver Operating Characteristic Curve (AUC-ROC) for classification [31], as well as Mean Squared Error (MSE), Root Mean Squared Error (RMSE), Mean Absolute Error (MAE), Coefficient of Determination (R²), and residual analyses for regression [31]. Model performance results are presented in an intuitive format that enables model comparison. Thus, users can assess model effectiveness and identify the most robust and predictive configurations with minimal manual intervention (see Fig. 1 and Supplementary Section 2.)

### Weighted Nested Cross-Validation for Unbiased Evaluation of Imbalanced Proteomic Data

Weighted nested cross-validation is a critical component of our workflow, designed to ensure unbiased model evaluation in the presence of extreme class imbalance, a common challenge in proteomic biomarker studies [12,32]. By organizing the process into an inner loop for hyperparameter optimization and an outer loop for performance evaluation, the nested framework effectively prevents information leakage and overfitting, thereby ensuring the generalizability of results [11]. To further mitigate the impact of unequal group sizes on model training and evaluation, we employ class weighting strategies throughout the workflow (see Supplementary Section 2). This includes integrating sample weights into the model fitting process, as well as computing weighted evaluation metrics, such as the weighted F1-score, which accounts for imbalanced class distributions by penalizing misclassifications proportionally to their class frequencies. This ensures that performance is not artificially inflated by dominant classes and reflects a model’s effectiveness across all groups. [19,33]. Additionally, we adopt stratified K-fold cross-validation to preserve the original class distribution within each fold, further enhancing the robustness and fairness of the evaluation [34]. When dimensionality reduction techniques are applied (e.g., PCA, UMAP), all feature selection steps are restricted to the inner loop, preserving the integrity of the validation process [11]. All reported evaluation metrics, including classification performance and regression error estimates, are derived strictly from the outer loop, providing reliable performance estimates suitable for high-dimensional, class-imbalanced proteomic data [11,12]. The structure of the weighted nested cross-validation scheme, incorporating stratified folds for both outer and inner resampling, is illustrated in Supplementary Information Section 2, Fig. S1.

### Integrative mean SHAP-Based Feature Interpretation with Network-Based Biomarker Expansion

To identify candidate biomarkers, mean SHAP values are used to compute feature importance in the best-performing model, estimating the contribution of individual proteins to model predictions [15,16]. One pitfall of interpreting ML/DL applications in biological systems is that ML/DL algorithms, as well as interpretative approaches such as SHAP, are designed to mitigate redundant information by selectively emphasizing one feature among pairs of highly correlated variables, whilst assigning lower importance to the other [15,16]. When a model favours one variable within a correlated set, the complementary variable may be assigned a lower mean SHAP value – not because it necessarily lacks relevance, but because its signal is already captured – resulting in an underestimation of its importance. However, in biological systems, particularly in molecular contexts such as protein co-expression and interaction networks, such correlations frequently reflect meaningful functional relationships and should not be dismissed as redundant noise [17,18]. Therefore, to reconcile the algorithmic handling of correlated variables in a biological setting, we identify an expanded list of putative biomarkers which themselves may have low mean SHAP values, but which are co-expressed with, and predicted interactors of, proteins with high mean SHAP values in the best performing model [35]. This additional step will enable researchers to conduct better downstream analyses of the biological processes and pathways underlying their studied disease, enabling a deeper understanding of disease mechanisms and potential therapeutic targets [36].

### Scalable, Cloud-Ready Deployment with Extensive Parallelization for Enhanced Computational Efficiency

Designed to meet the demands of large-scale proteomic studies, our workflow is implemented in WDL and is fully deployable on both cloud-based and high-performance computing platforms [37]. The system leverages advanced parallel computing strategies, including multi-threaded execution in frameworks such as XGBoost and LightGBM, parallelized hyperparameter tuning with Optuna [13,14], and concurrent processing of cross-validation folds across multiple CPU cores [38]. Furthermore, the computationally intensive SHAP calculations are optimized via parallel execution at both model and sample levels, thereby significantly reducing runtime without compromising accuracy [39].

### Report Generation

The workflow automatically generates a detailed PDF report that consolidates all major analytical outputs into a single, structured document. The report includes:

i. dimensionality reduction visualizations (e.g., PCA, UMAP) to aid in data exploration;
ii. comparative summaries of model performance across all trained ML and DL algorithms;
iii. per-model diagnostic visualizations, including confusion matrices, evaluation metric plots, mean SHAP value summaries, ROC curves, AUC and F1 score distributions across five cross-validation folds, as well as prediction-versus-observation plots and residual plots for both classification and regression tasks; and
iv. PPI network construction of top-ranked proteins, including first-degree network expansions to identify additional candidate biomarkers beyond those directly prioritized by SHAP.

This integrated reporting approach enhances both reproducibility and interpretability, facilitating downstream analysis and communication of results across interdisciplinary teams.

## Use case

### Proteomics Data for Classifying HBV-associated Liver Diseases

We applied BiomarkerML to reanalyze a publicly available urinary proteomic dataset originally published by Zhao et al. [40], generated as part of a study which sought to identify non-invasive biomarkers for hepatocellular carcinoma (HCC) in patients with chronic hepatitis B virus (HBV)-related liver diseases, including chronic hepatitis B (CHB) and liver cirrhosis (LC). From the original data, we selected 116 samples (23 CHB, 39 LC, and 54 HCC cases) with complete quantitative measurements for 2,868 urinary proteins. Detailed demographic information is presented in Table 1; full procedures for sample processing and proteomic analysis have been previously described [40].

**Table 1.** Demographics of populations in three groups.

| Disease | Number | Age | Sex (Male/Female) |
| --- | --- | --- | --- |
| CHB | 23 | 41.8 ± 10.9 | 17/6 |
| LC | 39 | 44.8 ± 8.3 | 31/8 |
| HCC | 54 | 54.93 ± 11.46 | 46/8 |

CHB = Chronic hepatitis B; LC= Liver cirrhosis; HCC = Hepatocellular carcinoma

In their original study, Zhao et al. implemented a two-stage biomarker discovery approach involving non-parametric Wilcoxon filtering followed by recursive feature elimination (RFE) and Random Forest classification, achieving an AUC of 0.871 in cross-validation. An ensemble voting classifier incorporating multiple base models further improved performance to an AUC of 0.903. A seven-protein panel was selected for modelling, and four of these proteins were subsequently validated in an independent cohort using a dataset that is not publicly available.

To ensure comparability, we first reproduced the binary classification setting of the original study (HCC vs. high-risk group [CHB + LC]) using the combined training and testing data. However, instead of applying univariate filtering or RFE, we employed feature selection and model training pipelines implemented within BiomarkerML. This is beneficial because the integrated pipeline enables automated, cross-validated feature selection in conjunction with model-specific hyperparameter tuning, thereby reducing the risk of information leakage and overfitting. Furthermore, it ensures consistency and reproducibility across models and datasets, while also facilitating SHAP-based interpretation of the selected proteins [11,13,15]. Then, to extend beyond the original binary setting, we applied BiomarkerML to the full three-class classification problem (CHB vs. LC vs. HCC), using the entire protein feature space without prior filtering.

### Binary Classification

#### Methods

The dataset described above was first converted into a standardized tabular format, where each row represented a sample and each column corresponded to a proteomic feature, with additional columns for sample identifiers and diagnostic labels. Samples were labelled as either high-risk (HR, comprising both LC and CHB) or HCC.

Binary classification was performed using a comprehensive set of candidate models, including RF, XGBoost, SVM, PLS-DA, MLP, KNN, Naïve Bayes, Logistic Regression, LightGBM, MLP within a VAE, and standalone VAE-based classification. The full set of proteins was retained for model training, without applying any prior dimensionality reduction.

All models were trained and evaluated using a nested cross-validation strategy to ensure robust and unbiased estimation of generalization performance, eliminating the need for manual train–test partitioning. Feature importance was assessed using mean SHAP values derived from the best-performing model.

#### Results

Following binary classification analysis using the BiomarkerML workflow, the performance metrics across all evaluated models were summarized (Fig. 2A). Among these, the LightGBM classifier exhibited the highest predictive performance, achieving an AUC of 0.93 (Fig. 2B). Model performance was consistent across outer cross-validation folds, supporting the robustness and generalizability of the LightGBM classifier (see Supplementary Section 3 and Supplementary Fig. S3).

**Figure 2.**
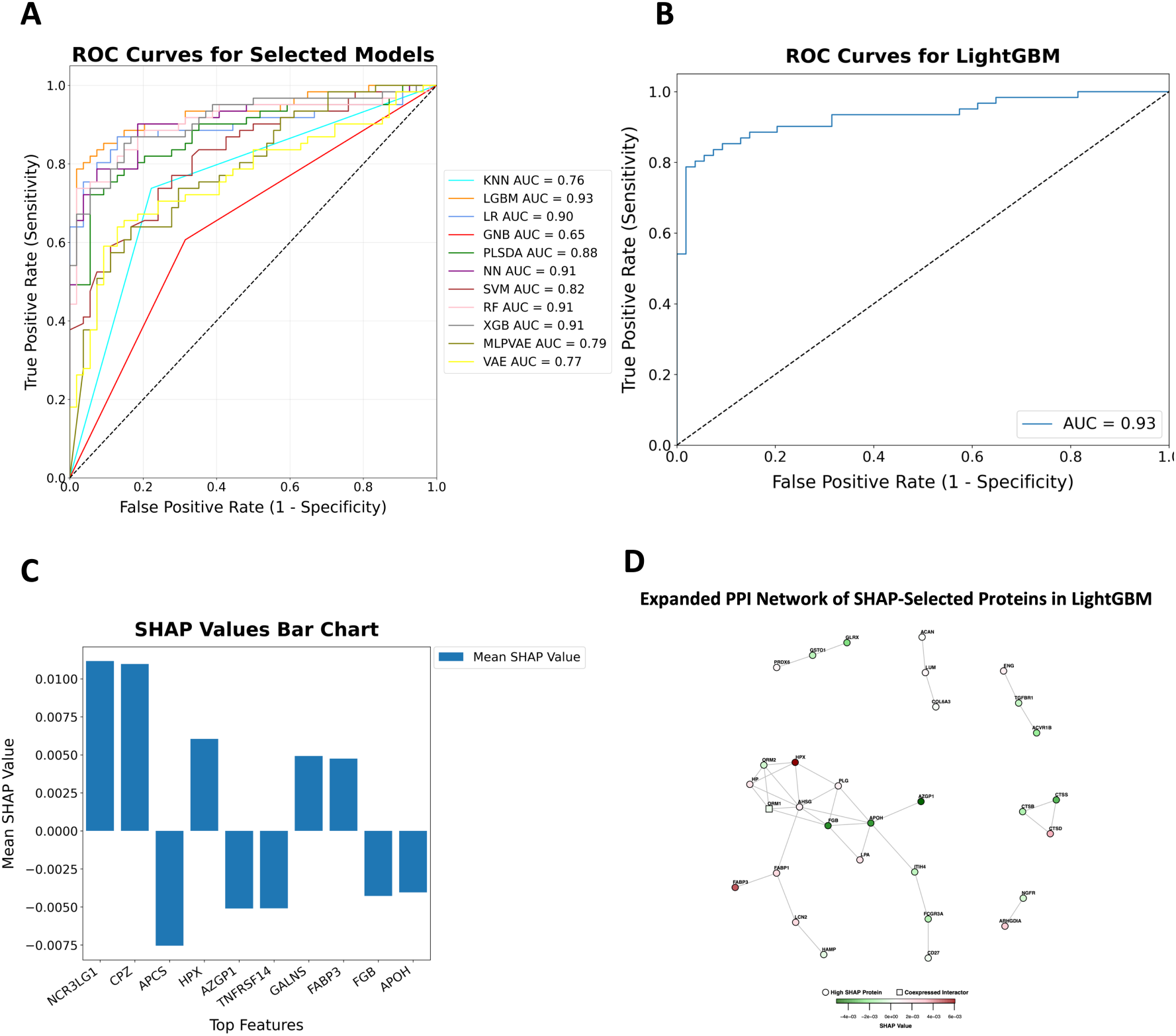
Performance evaluation of binary classification models. **(A)** AUC scores summarizing predictive performance across all evaluated models using nested cross-validation. **(B)** Receiver operating characteristic (ROC) curve for the best-performing model, LightGBM. **(C)** mean SHAP summary plot showing the top 10 proteins contributing to HCC vs. HR classification in the LightGBM model. **(D)** Expanded PPI network constructed from the top 100 proteins ranked by mean SHAP value using interaction data from the STRING database (combined score ≥ 800). The network was expanded to include co-expressed interactors of these proteins (combined score ≥ 800, Spearman’s ρ > 0.8). Top SHAP-ranked proteins are depicted as circles, and co-expressed interactors as squares. Node colour reflects the mean SHAP value in the LightGBM model.

To enhance interpretability, the top 10 proteins ranked by mean SHAP values from the LightGBM model are shown in Fig. 2C. These mean SHAP summary plots illustrate each feature’s contribution to the model’s predictions, with positive or negative mean SHAP values reflecting the direction and magnitude of influence on classification outcomes. An expanded PPI network constructed from the top 100 proteins ranked by mean SHAP value, with a one-degree expansion to include highly co-expressed interactors, is shown in Fig. 2D (see Supplementary Section 2 for details). Consistent performance across outer cross-validation folds indicates that the LightGBM classifier is both robust and generalizable (see Supplementary Section 3 and Supplementary Fig. S3C). Full evaluation metrics for LightGBM are provided in Supplementary Section 3 and Supplementary Fig. S3. Although our workflow supports the integration of dimensionality reduction techniques alongside machine learning models, incorporating ElasticNet prior to classification did not improve performance in this case (see Supplementary Section 3 and Supplementary Fig. S4).

### Multiclass Classification

#### Methods

As described above, the samples in the original data obtained from Zhao et al. [40] were labelled as CHB, LC or HCC. Global data structure and potential class separation was assessed using PCA, PLS, UMAP, and t-SNE for dimensionality reduction, followed by visualisation (Supplementary Section 4 and Figure S5). Then, multiclass classification was performed on the full data (without dimensionality reduction) across all candidate models, including RF, XGBoost, SVM, PLS-DA, MLP, KNN, Naïve Bayes, Logistic Regression, LightGBM, VAE and MLP in VAE. All models were trained and evaluated using nested cross-validation to ensure robust and unbiased performance estimation while avoiding manual train-test splits. Feature importance was interpreted via mean SHAP values computed from the best performing model.

#### Results

Following classification analysis using the BiomarkerML workflow, the averaged performance metrics for all evaluated models were summarized (Fig. 3A). Given the imbalance in class distribution across diagnostic categories (CHB, LC, HCC), we used macro-averaged AUC as the primary evaluation metric because this is the most appropriate metric for imbalanced multiclass problems (see Supplementary Section 2). LightGBM exhibited the strongest predictive performance, achieving a macro-averaged area under the receiver operating characteristic curve (AUC) of 0.84 (Fig. 3B). Model performance was consistent across outer cross-validation folds, supporting the robustness and generalizability of the LightGBM classifier (see Supplementary Section 4 and Supplementary Fig. S6C). Full model evaluation metrics for LightGBM are provided in Supplementary Section 4 and Supplementary Fig. S6.

**Figure 3:**
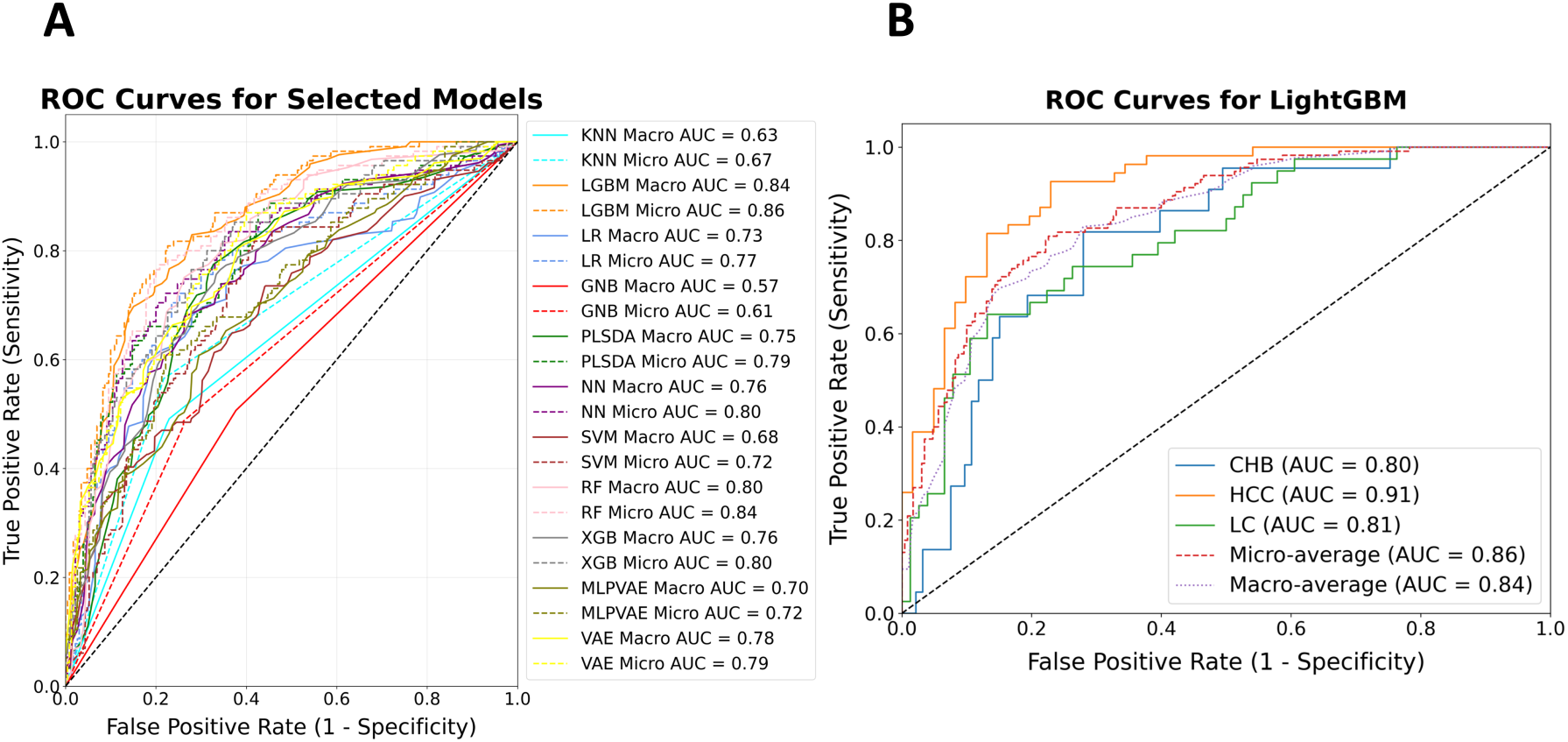
Performance evaluation of the classification models. **(A)** Macro-averaged AUC scores summarizing predictive performance across all evaluated models, based on nested cross-validation. **(B)** Receiver operating characteristic (ROC) curves for the LightGBM model, including both macro-averaged and class-specific curves (CHB, LC, HCC).

To facilitate model interpretability, proteins were ranked by the sum of their absolute mean SHAP values across all classes within the LightGBM model, and the top ten proteins were identified (Fig. 4A). Then, an expanded PPI network was constructed from the top 100 proteins, with a one-degree expansion to include directly connected co-expressed interactors (Fig. 4B; see Supplementary Section 2 for details).

**Figure 4.**
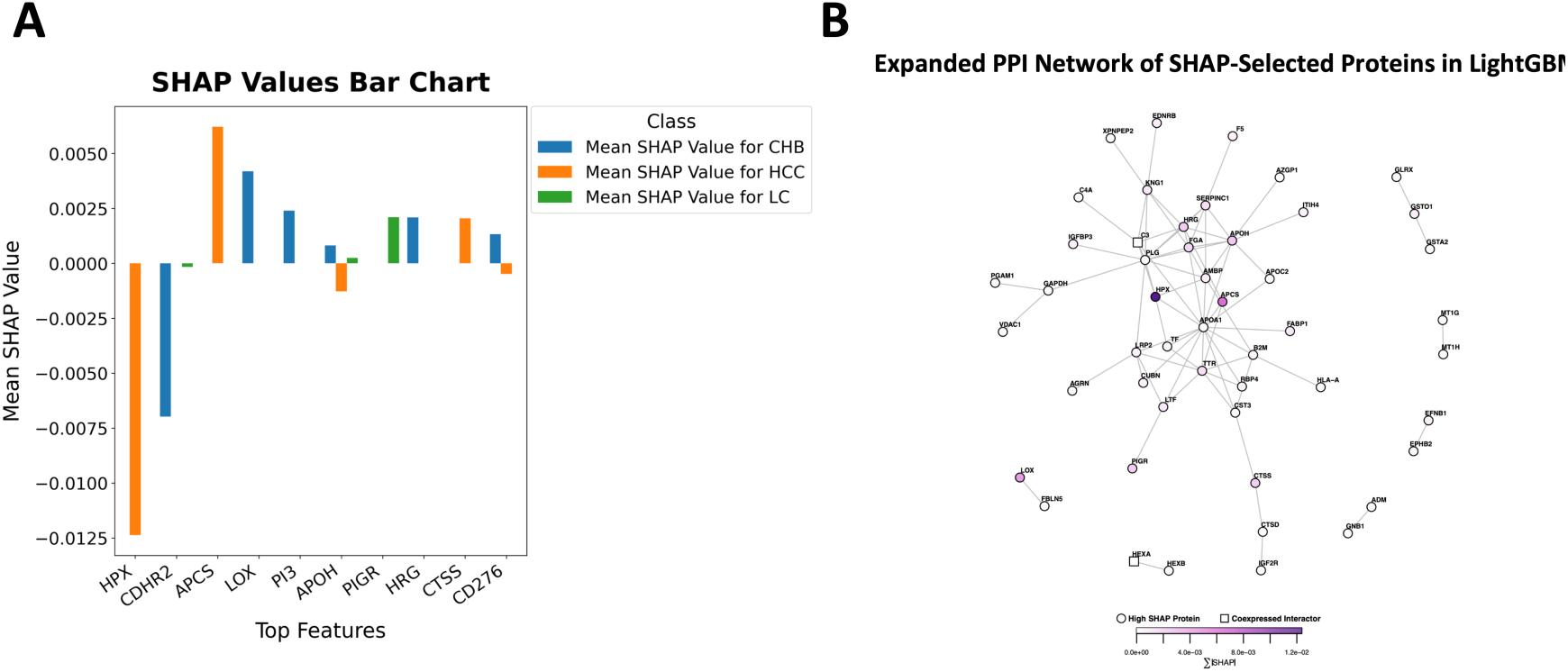
Interpretability and network-based contextualization of predictive proteins. **(A)** Mean SHAP summary plot showing the top 10 proteins contributing to LightGBM model predictions across all diagnostic classes (CHB, LC, HCC). Positive mean SHAP values indicate increased likelihood for a given class, while negative values indicate suppression. **(B)** Expanded PPI network constructed from the top 100 proteins ranked by the sum of their absolute mean SHAP values across all classes within the LightGBM model using interaction data from the STRING database (combined score ≥ 800). The network was expanded to include co-expressed interactors of these top 100 proteins (combined score ≥ 800, Spearman’s ρ > 0.8). Top SHAP-ranked proteins are depicted as circles, and co-expressed interactors as squares. Node colour reflects the mean SHAP value in the LightGBM model.

#### Discussion

In their original binary classification analysis distinguishing HCC from HR individuals, Zhao et al. [40] employed a two-stage feature selection strategy to identify urinary biomarkers distinguishing the two groups. Initially, candidate proteins were filtered using the Wilcoxon rank sum test (nominal p < 0.05, without multiple testing correction). Subsequently, recursive feature elimination (RFE) coupled with a Random Forest (RF) classifier was applied to refine feature selection and build predictive models. The resulting seven-protein panel achieved an AUC of 0.87 in the independent test dataset using the RF model, while a simpler voting-based ensemble classifier incorporating the same proteins yielded a higher AUC of 0.90, supporting its potential utility in clinical application. Of the seven evaluated proteins (HPX, APOH, PLG, APCS, TTR, SERPINC1, and GLRX), four (HPX, APOH, PLG, and GLRX) demonstrated the most consistent association with disease.

Here, we used BiomarkerML to perform a comparative analysis, but without the univariate pre-filtering step. In our analysis, BiomarkerML achieved an AUC of 0.93 using a single LightGBM model (Fig. 2B). Three biomarkers identified by Zhao et al., HPX, APCS, and APOH, were among the top 10 SHAP-ranked proteins (Fig. 2C), confirming both predictive and biological consistency across methods. In addition, GLRX and PLG ranked 17th and 47th, respectively, among the top SHAP-scoring proteins. Both were also located within the same connected subnetwork in the STRING-derived PPI network constructed from the top 100 SHAP-ranked proteins (Fig. 2D), further supporting their potential biological relevance. A list of ranked proteins along with their corresponding mean SHAP values is provided in the supplementary file: *Binary_Classification_lightgbm_shap_values.csv*.

In the multiclass setting, all seven proteins from Zhao et al. ranked within the top 5% of 2868 SHAP-scoring proteins. Among them, HPX, APCS, and APOH appeared within the top 10 ranked SHAP-scoring proteins (Fig. 4A). The remaining four proteins—PLG, TTR, GLRX and SERPINC1—were located within the same connected subnetwork in the STRING-derived expanded PPI network constructed from the top 100 SHAP-ranked proteins and their co-expressed interactors (Fig. 4B). Beyond these previously reported proteins, additional biomarker candidates with strong class-specific contributions emerged. For example, four of the top 10 SHAP-scoring proteins—HRG, LOX, PIGR, and CTSS— exhibited positive mean SHAP values primarily associated with distinct disease subtypes: HRG and LOX with CHB, PIGR with LC, and CTSS with HCC (Fig. 4A). All of these proteins also appeared in the same connected subnetwork as HPX, APCS, and APOH in the STRING-derived expanded PPI network (Fig. 4B). Beyond high SHAP-scoring proteins, two additional potentially biologically relevant proteins with low mean SHAP values were identified through PPI network expansion, namely, C3 and HEXA (Fig. 4B). A list of ranked proteins along with their corresponding mean SHAP values is provided in the supplementary file: *Multiclass_Classification_lightgbm_shap_values.csv*.

These observations demonstrate that our workflow replicates and improves on the predictive capacity of earlier methods; while Zhao et al.’s approach achieved strong binary performance, our workflow yielded improved AUCs (0.93 vs. 0.87–0.90) and maintained robust performance in the more complex multiclass setting (AUC 0.84). Furthermore, we have shown the capacity of BiomarkerML to validate biomarker robustness under class imbalance and multi-task settings, to enhance model interpretability using SHAP-based explanation methods, and to uncover additional putative biomarkers using expanded PPI networks. These results highlight the value of reproducible, interpretable, and extensible machine learning workflows in advancing biomarker discovery from complex proteomics datasets; in this case, ultimately supporting the development of reproducible and interpretable non-invasive biomarker strategies for HCC diagnosis.

## Implementation

BiomarkerML is written in the Python programming language and integrates several libraries for comprehensive data processing and analysis [41]. The *pandas* library and *NumPy* library are used for data storage and processing [42], while *scikit-learn* is employed for feature preprocessing, encoding, dimensionality reduction, and traditional model training [29]. All deep learning components, including the custom VAEs, are implemented in native *PyTorch* without reliance on third-party abstractions. Automated hyperparameter tuning is performed using the *Optuna* library within a nested cross-validation framework [13]. Model training is conducted using libraries such as *scikit-learn, XGBoost, LightGBM*, and *PyTorch* [43], and model evaluation-encompassing performance metrics and ROC curve analysis-is facilitated by *scikit-learn* [29]. Interpretability is enhanced through the *SHAP* library for computing feature importance [15], and data visualization is performed using the *seaborn, matplotlib*, and *plotly* libraries [44]. Finally, parallel computing is optimized using the *joblib* library, ensuring efficient execution during cross-validation, mean SHAP value computation, and hyperparameter tuning [45]. For a comprehensive description of the methodologies, please refer to Supplementary Section 2.

For workflow orchestration, BiomarkerML is written in Workflow Description Language (WDL) and run using the Cromwell engine. Each analytical step is defined as a modular task, enabling reproducibility, scalability, and seamless deployment across cloud-based and high-performance computing environments. All dependencies are containerized using Docker, ensuring full portability, version control, and long-term reproducibility. To run the workflow, users input a comma separated value (CSV) file formatted table of samples (rows) by protein expression values (columns). Additionally, users input metadata specifying dimensionality reduction strategies, classification or regression model choices, and selected analysis modules. Final outputs-including visualizations (.png), results tables (.csv), and model artifacts (.pkl, .npy) are consolidated into downloadable reports (.pdf, .tar.gz) for downstream interpretation. A schematic overview of the cloud-based proteomics machine learning workflow is provided in Supplementary Section 2 (Fig. S2). For full usage instructions see https://github.com/anand-imcm/proteomics-ML-workflow.

## Limitations

While BiomarkerML offers a powerful tool for proteomics analysis, it is important to acknowledge potential limitations. The effectiveness of the workflow depends on the quality of the input data, and any biases or errors in the initial data could impact the final outcomes. Further, although we set reasonable and conventional hyperparameter ranges and utilize *Optuna* with cross-validation to automatically find optimal settings, interaction with the model may still be required to fine-tune results in cases where the data presents unique challenges (e.g., highly imbalanced classes, sparsity, or non-standard distributions). Additionally, while the workflow includes a wide range of models and techniques, future work could focus on expanding this repertoire to include emerging methods and addressing potential biases in the data, such as the missing data imputation process [46,47].

## Conclusion

BiomarkerML represents a substantial methodological advancement in proteomic data analysis, offering an integrated, scalable, and reproducible workflow that addresses key challenges inherent to proteomic datasets—including non-linearity, high dimensionality, and biological complexity. By systematically combining advanced machine learning and deep learning techniques with robust dimensionality reduction, model selection, SHAP-based feature importance evaluation, and expanded PPI network construction, the framework enables reliable identification of diagnostically relevant protein biomarkers and meaningful biological insights.

Designed with flexibility and extensibility in mind, BiomarkerML supports automation of complex analytical steps and is accessible to researchers without deep computational expertise. All components, including two custom-built generative AI architectures, are implemented in native Python, *Scikit-learn* and *PyTorch* code, ensuring full transparency and code-level reproducibility. The incorporation of parallel computing further enhances performance for large-scale data.

Overall, BiomarkerML enables researchers to rapidly identify interpretable and biologically grounded candidate biomarkers from high-throughput proteomic data, so it is well positioned to serve as a foundational tool in systems biology and precision medicine, adaptable to evolving technologies and future research needs.

## Outlook

Despite the aforementioned limitations, several promising avenues exist for the future development of BiomarkerML. A key objective is to enhance its capabilities by integrating advanced deep learning architectures, particularly for longitudinal data analysis and multi-modal datasets, where complex temporal dependencies and heterogeneous data sources pose significant analytical challenges. Expanding BiomarkerML in this manner will not only improve its adaptability to dynamic and high-dimensional biological data but also broaden its applicability across a diverse range of omics studies, biomarker discovery, and drug development pipelines. By incorporating more sophisticated deep learning models, BiomarkerML aims to facilitate predictive modelling, patient stratification, and mechanistic insights, thereby strengthening its role as a powerful tool for precision medicine and translational research.

## Availability of supporting source code and requirements

List the following:

- Project name: BiomarkerML
- Project home page: https://github.com/anand-imcm/proteomics-ML-workflow
- Operating system(s): Linux (64-bit)
- Programming language: Python, R and WDL
- Other requirements: Docker and Cromwell
- License: BSD-3-Clause
- RRID: BiomarkerML (RRID:SCR_027268)

## Additional files

SupplementaryTableS1.xlsx

## Code availability

The source code, including the usage instructions can be found at https://github.com/anand-imcm/proteomics-ML-workflow

## Abbreviations

ML: machine learning
DL: deep learning
WDL: Workflow Description Language
SHAP: SHapley Additive exPlanations
MS: Mass spectrometry
RF: Random Forest
SVM: Support Vector Machine
VAE: Variational Autoencoders
MLP: Multilayer Perceptron
XGBoost: Extreme Gradient Boosting
AUC: Area Under the Curve
ROC: Receiver Operating Characteristic Curve
MSE: Mean Squared Error
RMSE: Root Mean Squared Error
MAE: Mean Absolute Error
R²: Coefficient of Determination
PCA: Principal Component Analysis
KPCA: Kernel Principal Component Analysis
t-SNE: t-distributed Stochastic Neighbor Embedding
PLS: Partial Least Squares
UMAP: Uniform Manifold Approximation and Projection
PPI: protein-protein interaction
CHB: Chronic hepatitis B
LC: Liver cirrhosis
HCC: Hepatocellular carcinoma
HR: high-risk

## Acknowledgements

The authors would like to thank Prerak Desai (GlaxoSmithKline) for helpful insights and discussions.

## Author’s contributions

AT conceived BiomarkerML and AT, YZ, AM, and YD collaborated to mature its design. YZ wrote the machine learning source code and conducted testing and debugging of all machine learning scripts. YD wrote the protein network analysis code. AM implemented the WDL workflow and conducted testing and debugging of BiomarkerML on our cloud platform testing environment. CR acquired use-case datasets from public databases and made them available on our cloud platform testing environment. YZ, AT, and MF wrote, proof-read and edited the bulk of the manuscript, with additional contributions from AM and YD pertaining to methodological details.

## Supplementary Information

### Section 1 Existing ML workflows for proteomic biomarker discovery

Attached Supplementary Table 1: Comparison of existing ML workflows for proteomic biomarker discovery

**Table S1:** While some existing workflows employ basic ML models, they are often limited in scope and rarely support advanced deep learning methods. For example, recent studies utilizing ML pipelines to analyze clinical and proteomic data have often employed a limited range of models, with ensemble voting used to select the best-performing model, rather than conducting a thorough comparison of individual models [1]. The AutoXAI4Omics workflow integrates standard ML classification and regression methods, facilitating model comparisons and interpreting features through SHAP values; however, it lacks more sophisticated feature selection preprocessing techniques and the incorporation of deep learning models [2]. Similarly, the AITeQ workflow includes ML models similar to those in AutoXAI4Omics but relies on differential analysis for preprocessing and does not perform model comparisons or identify key features [3]. The MLme workflow incorporates feature selection and standard ML classification models, generating reports to clarify results; however, its pipeline is incomplete, lacking hyperparameter fine-tuning, model comparison, feature importance analysis, and subsequent interpretation of results [4]. The HiOmics workflow offers a more comprehensive cloud-based pipeline that includes standard ML modeling, result comparison, and feature importance assessment, as well as downstream bioinformatics analyses and report generation [5]. However, despite its broader scope, HiOmics is still limited by the absence of feature selection, regression analysis, and deep learning models, and is not specifically tailored for proteomics data [5]. The MLOmics framework integrates a diverse set of literature-derived machine learning and deep learning models into a standardized evaluation pipeline for cancer multi-omics data, supporting classification, clustering, and imputation tasks [6]. However, it does not include hyperparameter optimization, regression analysis, or feature interpretability methods such as SHAP. In addition, its implementation centres on integrating previously published models, rather than providing a modular framework for custom model development or task-specific adaptation [6].

### Section 2 BiomarkerML implementation details

#### Data Preprocessing

##### Data standardization

The dataset loads in CSV format, with the column labeled “Label” containing sample annotations, and the column “SampleID” including Samples’ ID. Samples with any *NaN* values are excluded to maintain data integrity. Features are standardized using *StandardScaler* from *sklearn. Preprocessing* in the weighted nested cross-validation, normalizing them to have a mean of zero and a standard deviation of one. The target variable is encoded into numerical labels using *LabelEncoder*. For multiclass classification tasks, labels are binarized using *pd.get_dummies* to facilitate the calculation of metrics such as ROC curves and AUC scores.

##### Dimensionality Reduction Techniques

To address the curse of dimensionality and enhance model performance, six dimensionality reduction techniques are applied [7]. The methodology integrates optional dimensionality reduction techniques and dataset-wide visualization to enhance data interpretability. Furthermore, feature selection is systematically embedded within the weighted nested cross-validation framework to optimize the performance of machine learning models. Features are standardized using *StandardScaler* prior to applying these techniques. A random state is set to ensure reproducibility.

1. Principal Component Analysis (PCA): PCA is conducted using the PCA class from *scikit-learn*, with the number of components optimized within a range of 2 to the maximum specified dimensions using *GridSearchCV*.
2. Uniform Manifold Approximation and Projection (UMAP): UMAP is implemented using the *umap-learn* library. The number of neighbours (n_neighbours) and the minimum distance (min_dist) were optimized using a variance-maximizing objective function within the ranges of 5–50 and 0.1–0.9, respectively. Parameter selection was performed using the Optuna framework to enhance separation in the low-dimensional space.
3. t-distributed Stochastic Neighbour Embedding (t-SNE): t-SNE is performed using the *TSNE* class from *scikit-learn*, with perplexity varied within a specified range. The configuration is optimized based on the highest variance observed in the transformed data.
4. Kernel Principal Component Analysis (KPCA): KPCA is conducted using the *KernelPCA* class from *scikit-learn*. Various kernel functions and the gamma parameter are explored, with the optimal combination selected based on maximizing the variance of the transformed components.
5. Partial Least Squares (PLS) Regression: PLS regression is performed using the *PLSRegression* class from *scikit-learn*, with the number of components optimized to minimize mean squared error. Labels are encoded into numeric form using *LabelEncoder*. The corresponding value for ‘Label’ needs to be provided for this method.
6. ElasticNet Regularization: ElasticNet regularization is applied using the *ElasticNetCV* class from *scikit-learn*, combining L1 and L2 penalties. The ratio of penalties is optimized across a specified range using 5-fold cross-validation. The loss function is based on least square loss. Features are selected based on the absolute magnitude of the Elastic Net coefficients and then standardized using *StandardScaler*. The corresponding value for ‘Label’ needs to be provided for this method.

##### Visualization and Results

Dimensionality reduction results are visualized using *seaborn’s PairGrid* functionality. Pairwise scatterplots and diagonal density plots are generated for each method, illustrating data structure in reduced-dimensional space. ElasticNet coefficients are visualized to highlight significant contributions. If the dimensionality reduction methods implement in advance, the workflow will calculate the transformed components’ SHAP values.

##### Classical Machine Learning Model Selection and Hyperparameter Tuning

A comprehensive set of classical supervised machine learning classification methods are employed with cross validation [8], using the *scikit-learn* and *xgboost* libraries. All classification models in BiomarkerML support both binary and multiclass classification tasks. The hyperparameters are tuned using *Optuna* [9,10], a Bayesian method-based hyperparameter optimization framework for ML/DL models in the weighted nested cross-validation. A random state is set to ensure reproducibility.

##### Machine Learning Model Details

1. Neural Network (MLP): The MLP classifier from *sklearn.neural_network.MLPClassifier*, is configured with a *ReLU* activation function and a maximum number of iterations. Hyperparameter tuning involves varying the number of hidden layers, the L2 regularization term, and the initial learning rate within specified ranges.
2. Random Forest(RF): The Random Forest model from *sklearn.ensemble.RandomForestClassifier*, is tuned by adjusting the number of trees, feature selection strategies, and tree depths.
3. Support Vector Machine (SVM): The SVM classifier from *sklearn.svm.SVC*, is implemented with a radial basis function (RBF) kernel, optimized for non-linear classification problems. Hyperparameter optimization focuses on adjusting the regularization parameter and the kernel coefficient.
4. Partial Least Squares Discriminant Analysis (PLSDA): PLSDA is implemented using *sklearn.cross_decomposition.PLSRegression*, serving simultaneously for dimensionality reduction and supervised classification. In binary classification, decision values are thresholded at 0.5; for multiclass problems, the class corresponding to the maximal predicted probability is assigned.
5. XGBoost: The XGBoost model from *xgboost.XGBClassifier*, is configured with the mlogloss evaluation metric for multiclass classification. Hyperparameter tuning involves varying the number of boosting rounds, tree depth, and learning rate.
6. K-Nearest Neighbors (KNN): The KNN model from *sklearn.neighbors.KNeighborsClassifier* is optimized by varying the number of neighbors. Weighted nested cross-validation framework is applied, where the inner loop selects the best number of neighbors through Optuna.
7. LightGBM: The LightGBM model from *lightgbm.LGBMClassifier*, is optimized using Optuna across several hyperparameters, including the number of boosting rounds, tree depth, and learning rate.
8. Naive Bayes: The Gaussian Naive Bayes classifier from *sklearn.naive_bayes.GaussianNB*, is used with default parameters, learning class priors from the data.
9. Logistic Regression: The Logistic Regression model *sklearn.linear_model.LogisticRegression*, is optimized by tuning the regularization strength within a specified range.

##### Deep Learning Model Selection and Hyperparameter Tuning

The two generative AI models that we contribute are based on *PyTorch* implementations, with customized architectural designs tailored to our task [11]. While our model builds upon the standard *PyTorch* implementation of the Variational Autoencoder (VAE), we have designed a customized architecture that integrates a tailored encoder-decoder structure and a downstream MLP classifier, trained jointly via a two-stage process with early stopping and *Optuna*-based hyperparameter optimization. This constitutes a key methodological contribution of our work [9,10]. A random state is set to ensure reproducibility.

1. Variational Autoencoder with Multilayer Perceptron (VAE-MLP): The VAE-MLP model implemented in *PyTorch* classifies the dataset by using the reconstructed data from the VAE as input to the MLP classifier. The VAE includes an encoder, a latent space, and a decoder, where the encoder transforms the input into a low-dimensional latent space, and the decoder reconstructs the input data. The MLP classifier, composed of fully connected layers with *LeakyReLU* activations, uses this reconstructed data for classification.
2. Multilayer Perceptron inside Variational Autoencoder (MLP in VAE) The second VAE-MLP model from *PyTorch* uses the encoded latent space representation (*z_mean*) from the VAE as the input for the MLP classifier. The VAE consists of an encoder, which projects the input into a low-dimensional latent space, and a decoder, but only the latent space representation (*z_mean*) is passed to the MLP for classification. The MLP, composed of dense layers with *LeakyReLU* activations, uses these latent space embeddings to predict the target classes.

##### Classification Model Evaluation and Metrics

Model performance is evaluated using several key metrics provided by *scikit-learn* [12]. Accuracy is calculated using *accuracy_score*, representing the proportion of correctly classified samples. The weighted F1 score, computed with *f1_score*, is used to balance precision and recall, accounting for class imbalance in multiclass scenarios. Confusion matrices are generated using *confusion_matrix* and visualized with *ConfusionMatrixDisplay*, providing insight into the distribution of true positives, false positives, true negatives, and false negatives for each class. Sensitivity (recall) is derived as the proportion of true positives out of the sum of true positives and false negatives, while specificity measures the proportion of true negatives out of all negative cases. These metrics help to evaluate both class-wise performance and the overall effectiveness of the models. Visualization of performance metrics, including bar charts, is done using *matplotlib*. The final model was selected based on the average performance across a weighted nested 5-fold stratified cross-validation scheme, using class-balanced sampling (*class_weight=’balanced’* or *sample_weight* as applicable) [13]. The outer loop evaluated generalization performance, while the inner loop optimized hyperparameters using training data only. Stratification ensured that each fold preserved the original class distribution, which is particularly important given the pronounced class imbalance in our dataset. To achieve this, we implemented *StratifiedKFold* from the *sklearn.model_selection* module, which guarantees that each fold contains approximately the same proportion of samples from each class, thereby preventing biased learning and preserving representativeness across splits. A weighted nested cross-validation scheme with stratified folds was employed for model training and evaluation (Figure S1 from [14,15]).

**Figure S1:**
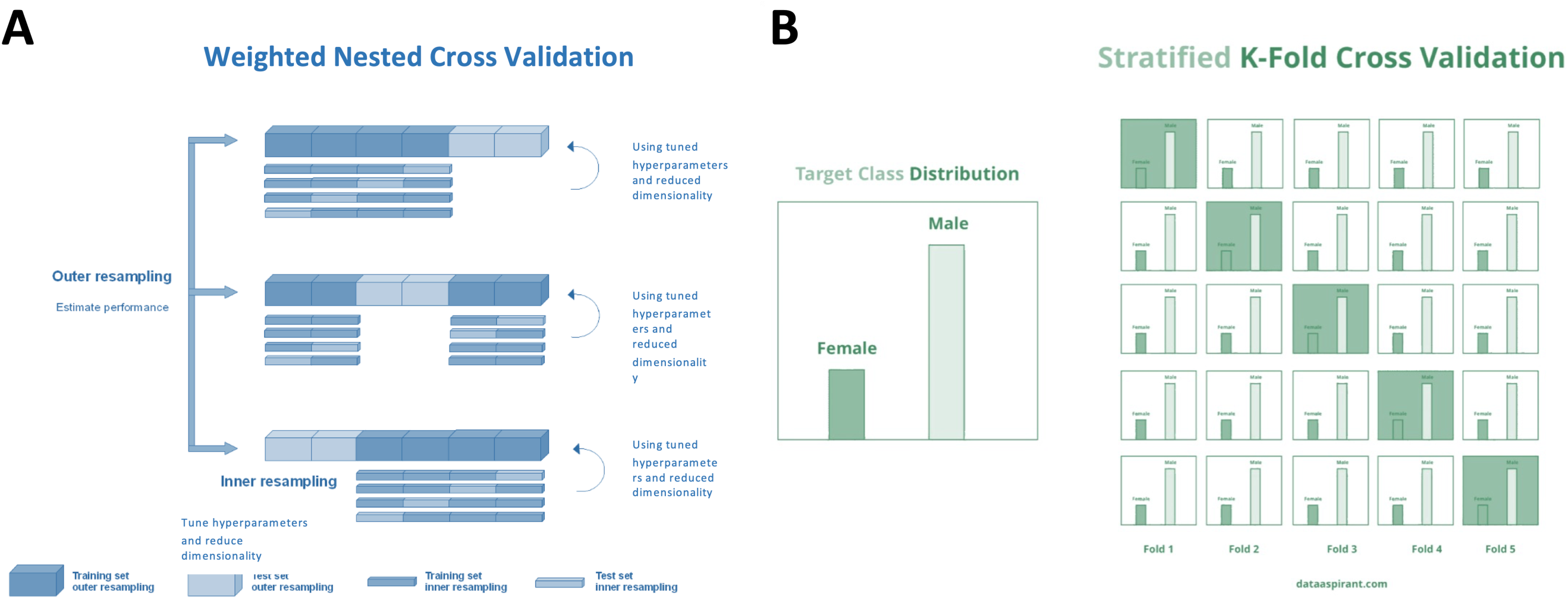
Schematic illustration of the cross-validation strategies used in BiomarkerML. A: Weighted nested cross-validation framework. An outer loop performs repeated stratified sampling to estimate generalization performance, while the inner loop tunes hyperparameters and selects features using stratified resampling within each outer fold. Class imbalance is accounted for by weighting the objective function or sampling procedure. **B: Stratified K-fold** cross-validation ensures that the distribution of classes (e.g., male vs female) is preserved across all training and validation splits [14,15].

##### ROC Curve Comparison Across Models

To compare the performance of all models, ROC curves are generated and plotted on a single graph using a custom script that utilizes *scikit-learn* and *matplotlib* [12]. The script loads ROC data for each selected model and uses scikit-learn’s *roc_curve* function to compute the false positive rates (FPR) and true positive rates (TPR), while the *auc* function calculates the area under the ROC curve (AUC). For binary classification models, the ROC curve for the single class is plotted, while for multiclass models, the micro-averaged ROC curve aggregates the true positive rates and false positive rates across all classes, and the macro-averaged ROC curve computes the ROC curve for each class independently and then averages the true positive rates and false positive rates across all classes. Unlike macro-averaging, which treats each class equally regardless of its frequency, micro-averaging aggregates predictions over all classes, thereby providing a performance estimate that is weighted by the actual sample distribution. This makes macro-averaged AUC more appropriate for imbalanced multi-class problems, such as those encountered in proteomics-based clinical classification. The *matplotlib* package is used to visualize the ROC curves and add legends displaying the AUC values for each model. Higher AUC values indicate better discriminatory power, making these models more effective at distinguishing between different classes.

##### Classical Machine Learning Regression Methods

A diverse set of classical supervised machine learning regression methods is employed by *scikit-learn* [11]. Each regression model is fine-tuned using *Optuna* to select the best-performing hyperparameters, followed by an evaluation using weighted nested 5-fold cross-validation to ensure robust performance. A random state is set to ensure reproducibility.

###### Regression Model Details

1. Neural Network (MLP): The MLPRegressor was configured with a maximum of 200,000 training iterations to ensure full convergence. Hyperparameter tuning involves varying the number of hidden layers and the L2 regularization term within specified ranges.
2. Random Forest: The *RandomForestRegressor* is tuned by adjusting the number of trees, feature selection strategies, and tree depths.
3. Support Vector Regression (SVR): The *SVR* is implemented with a function kernel. Hyperparameter optimization focuses on adjusting the regularization parameter and the kernel coefficient.
4. XGBoost: The *XGBoostRegressor* is configured with the *rmse* evaluation metric for regression tasks. Hyperparameter tuning involves varying the number of boosting rounds, tree depth, and learning rate.
5. Partial Least Squares Regression (PLS): The *PLSRegression* model is optimized by varying the number of components within the model.
6. K-Nearest Neighbors Regression (KNN): The *KNeighborsRegressor* is optimized by varying the number of neighbors.
7. LightGBM: The *LightGBMRegressor* is optimized using a Optuna across several hyperparameters.

##### Regression Model Evaluation and Metrics

The performance of each regression model is evaluated by the average performance using weighted nested 5-fold cross-validation. Evaluation metrics include Mean Squared Error (MSE), Root Mean Squared Error (RMSE), Mean Absolute Error (MAE), and R-squared (R^2^) scores, using *sklearn.metrics [12]*. Predicted values are plotted against actual values to generate prediction graphs, and residuals are plotted to assess model fit. All models are saved along with their parameters, results, and associated plots for further analysis. The hyperparameters are tuned using *Optuna* [10].

##### SHAP Analysis

To interpret the contribution of each feature to the model’s predictions, SHAP (SHapley Additive exPlanations) values are computed for each model using a model-specific SHAP explainer [16].

For tree-based models, such as Random Forest, XGBoost, and LightGBM, *shap.TreeExplainer* is employed, leveraging its optimization for tree-structured algorithms. For other models, including Neural Networks, PLSDA, Logistic Regression, KNN, SVM, and deep learning architectures like VAE-MLP, the model-agnostic *shap.PermutationExplainer* is used. If dimensionality reduction is applied prior to model fitting, SHAP values are computed for the resulting transformed components (e.g., principal components), rather than the original input features.

SHAP values are visualized using different methods depending on the implementation: radar charts are used to highlight top features with the highest mean SHAP values, while bar plots and summary dot plots are generated to show the most influential features contributing to model predictions. To summarize feature-level importance, SHAP values are averaged across all samples. In binary classification, this results in a single mean SHAP value per feature; in multiclass classification, one mean SHAP value is computed per feature per class. These are referred to throughout the manuscript as “mean SHAP values.” Selection of top-ranked proteins was based on the absolute value of their mean SHAP scores. For multiclass classification tasks, selection was performed using the sum of absolute mean SHAP values across all classes for each protein.

##### Parallel Computing

To optimize the performance of machine learning models, we implement parallel computing techniques to reduce computation time, particularly during cross-validation and hyperparameter optimization [17,18]. Models such as XGBoost, LightGBM, SVM, and Neural Networks are configured using the *cross_val_predict* function with the *n_jobs=-1* argument, allowing parallel execution across all available CPU cores. This facilitates the simultaneous evaluation of multiple cross-validation folds, significantly enhancing computational efficiency. Libraries like XGBoost and LightGBM, which natively support multi-threading, further accelerate the model fitting process by distributing tasks such as tree construction and gradient boosting across multiple threads. Parallel computation is also implemented using *joblib.Parallel* to speed up the model training and evaluation during cross-validation. For each fold, the *Parallel* function, combined with the function parameter *delayed*, independently trains and evaluates the model for each fold, thus parallelizing the entire cross-validation process.

For hyperparameter optimization, we employ *Optuna*, which supports parallelized evaluation of hyperparameter combinations. This setup reduces the time required for each trial by distributing computations across multiple cores. Additionally, SHAP value calculations are integrated into the workflow using *ProcessPoolExecutor* and *joblib.Parallel* to enable parallel execution across multiple models. This approach allows SHAP computations to leverage multi-core CPU environments, particularly those with high-core processors, significantly enhancing computational efficiency. Furthermore, SHAP’s *PermutationExplainer* is parallelized at the sample level via the *n_jobs_explainer* parameter, facilitating the simultaneous computation of SHAP values for multiple samples. For multi-class classification tasks, *joblib.Parallel* computes SHAP values across different classes concurrently, optimizing the process and ensuring high efficiency across complex classification models.

##### Protein-Protein Interaction Network and Expanded Protein-Protein Interaction Network

To interpret the biological relevance of model-derived features, we constructed protein–protein interaction (PPI) networks using the top-ranked proteins based on their mean SHAP values (as described in subsection SHAP Analysis, above). PPI networks were derived from the STRING database (version 12.0, human, taxon ID 9606), which assigns each potential interaction a combined score (0–1000) that integrates multiple lines of evidence, including experimental data, curated databases, text mining, gene co-expression, and computational predictions. To control the inclusion of protein–protein edges, an evidence score threshold was applied. In BiomarkerML. this threshold is user-defined (via the *--combined_score_thresholdHere* option), with a default of combined score ≥ 800, thereby allowing flexibility in defining interaction stringency and focusing on biologically meaningful connections.

Given that machine learning models inherently account for feature collinearity, some biologically relevant proteins that are highly co-expressed with high mean SHAP value proteins may receive lower mean SHAP values themselves. To address this, our pipeline supports the construction of expanded PPI networks by identifying proteins that are both high-confidence interactors (based on the combined score threshold defined above) and strongly co-expressed (Spearman’s ρ > 0.8) with top-ranked SHAP proteins.

##### PDF Report Generation

To facilitate comprehensive analysis and easy dissemination of results, a PDF report is automatically generated, summarizing the outcomes of the model evaluations. The report includes sections such as an introduction, detailed analysis of model performance, and visualizations of the results. For each machine learning model, the report presents confusion matrices, evaluation metrics and SHAP radar plots, alongside visualizations from various dimensionality reduction techniques. This report serves as a convenient reference, consolidating the analysis into a single accessible document.

##### WDL wrappings for workflow

The main workflow is implemented in WDL v1.0 (Workflow Description Language) and can be executed using the *Cromwell* engine. The workflow has a modular design, where every step of the analysis is defined into standalone *tasks*. These tasks consist of the declaration of the inputs, outputs and runtime environment. The modules for dimensionality reduction, classification, regression, SHAP calculation, protein network analysis and summarization are written into individual *.wdl* files. This allows easy maintenance of the workflow and flexibility to be able to update or add new modules.

The modules are then called using the *main.wdl* file as per the user’s input. The tasks for dimensionality reduction, classification and regression are executed in parallel using WDL’s *scatter-gather* functionality based on the user’s input.

The dependencies are managed using Docker and all the WDL tasks are configured to run in Docker containers to ensure seamless portability across different computing environments. A schematic overview of the cloud-based proteomics machine learning workflow is provided in Figure S2.

**Figure S2:**
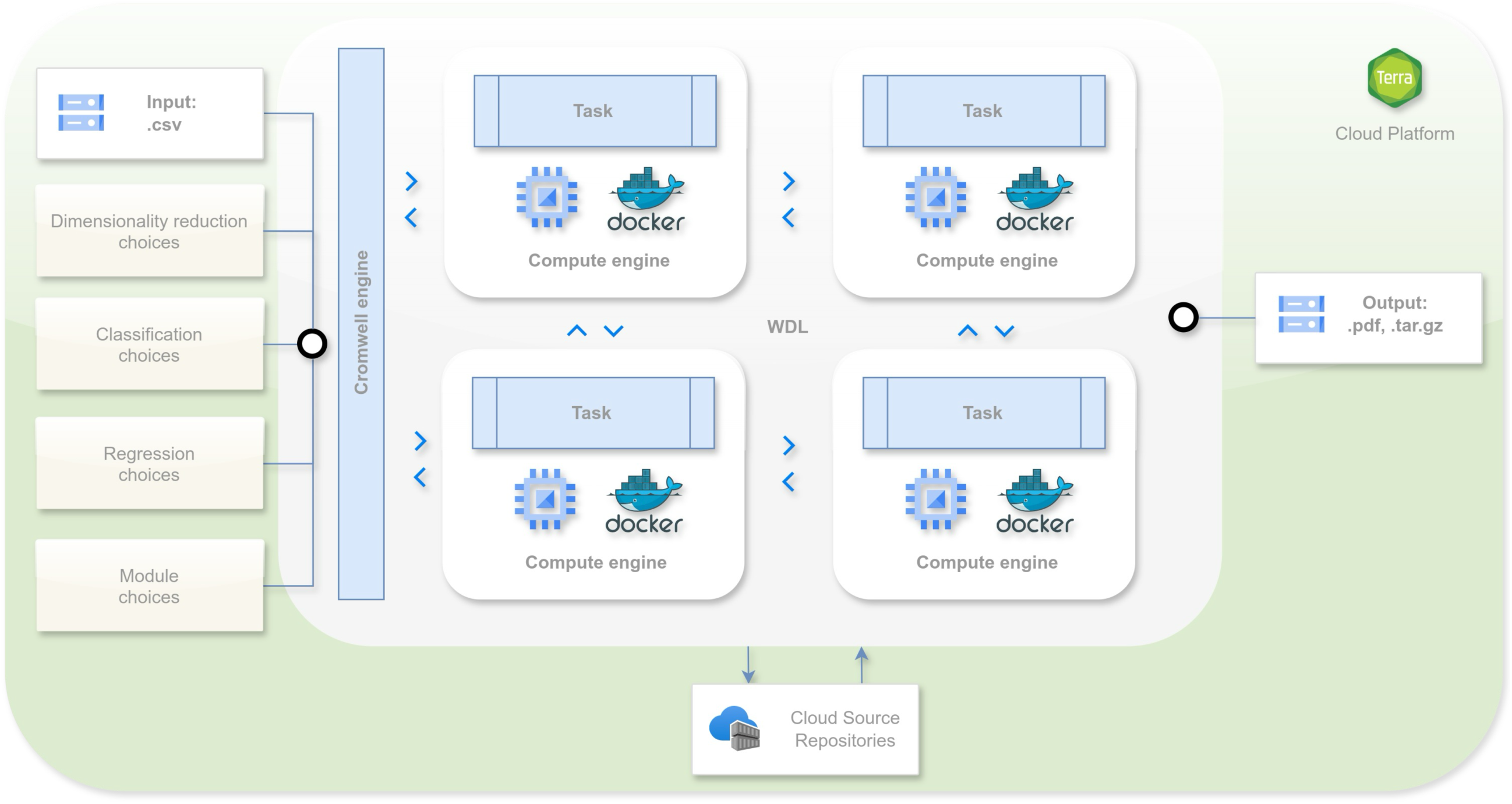
Graphical overview of the cloud-based proteomics machine learning workflow. The pipeline takes as input a user-provided CSV dataset (samples × features) along with metadata specifying dimensionality reduction strategies, classification or regression model choices, and selected analysis modules. Tasks are orchestrated by the Cromwell engine using WDL scripts and executed in parallel via Docker-enabled compute engines. Final outputs-including visualizations (.png), results tables (.csv), and model artifacts (.pkl, .npy)-are consolidated into downloadable reports (.pdf, .tar.gz) for downstream interpretation.

### Section 3 Full Results of Binary Classification Analysis

#### Full Results of Binary Classification Analysis - without ElasticNet

**Figure S3:**
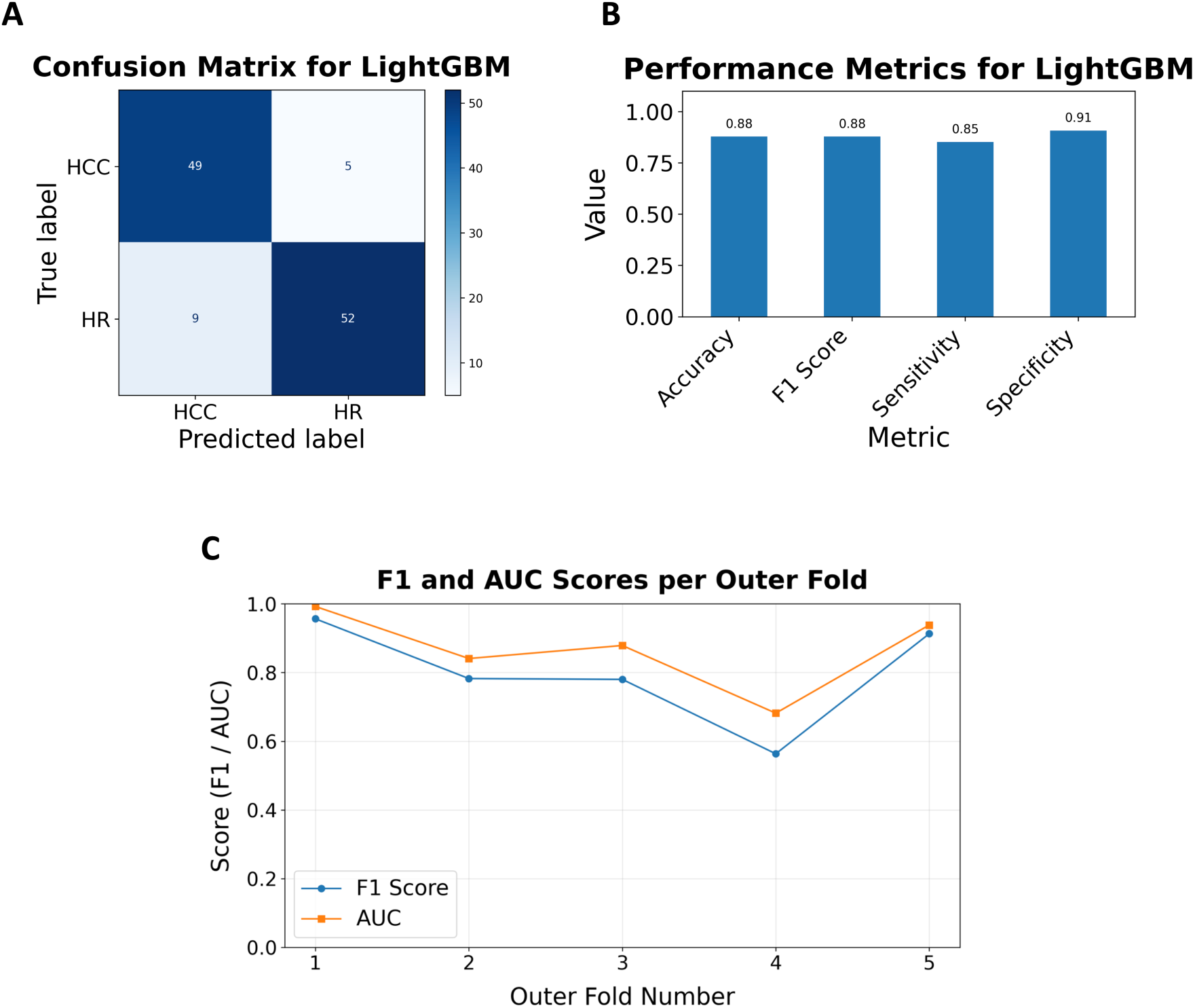
Performance evaluation and model interpretability for binary classification tasks in liver disease. **(A)** Confusion matrices depicting class-level prediction performance for LightGBM. **(B)** The evaluation of accuracy, sensitivity, specificity, and weighted F1-score for the model. **(C)** Outer cross-validation results showing fold-wise distributions of AUC and F1-score, illustrating model stability and generalization.

#### Full Results of Binary Classification Analysis - with ElasticNet

**Figure S4:**
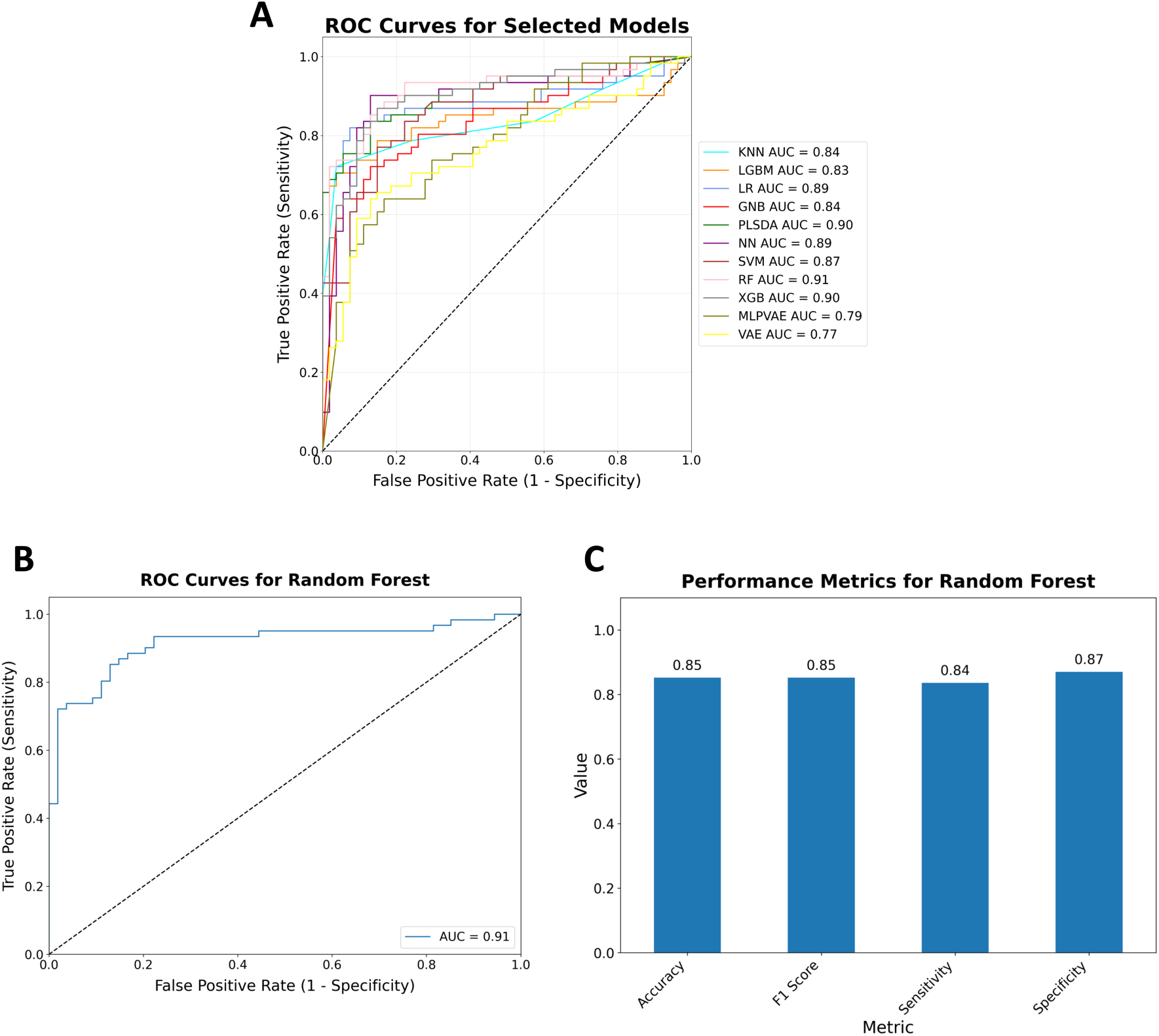
Performance evaluation and model interpretability for binary classification tasks in liver disease. **(A)** Area under the ROC curve (AUC) comparison across all evaluated classifiers. **(B)** Receiver operating characteristic (ROC) curves for the top-performing models: Random Forest (RF). **(C)** The evaluation of accuracy, sensitivity, specificity, and weighted F1-score for the model.

### Section 4 Full Results of Multiclass Classification Analysis

#### Global structure visualization using dimensionality reduction

BiomarkerML provides methods for visualising dimensionally reduced data to assess global data structure prior to modelling. Here, we used BiomarkerML to project sample distributions onto three dimensions using PCA, PLS, UMAP, and t-SNE, followed by visualisation using scatter plots and histograms (Fig S5 A-D). These visualizations provide a practical approach for preliminary quality assessment prior to running computationally intensive modelling workflows.

**Figure S5:**
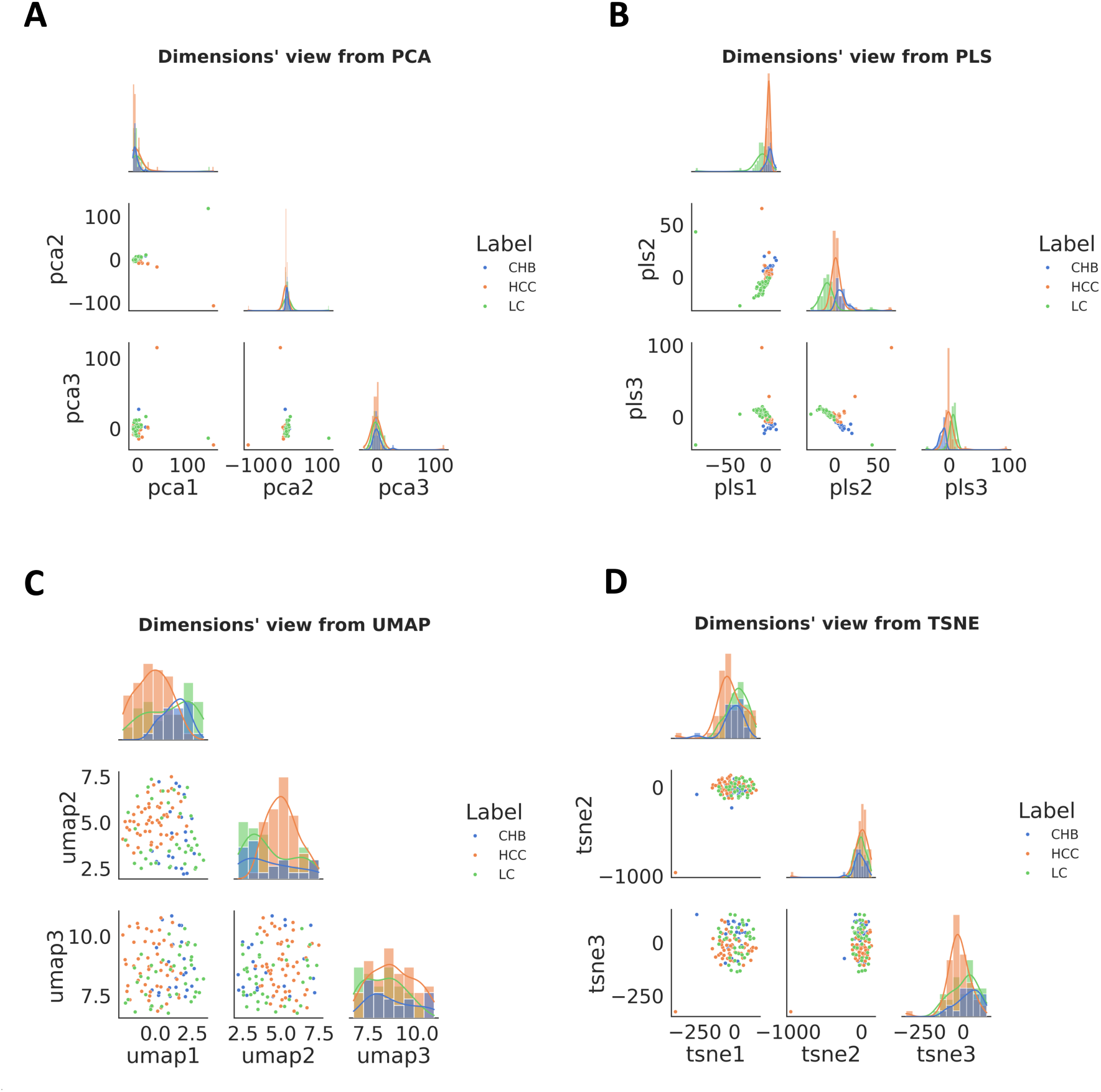
Dimensionality reduction and visualization of proteomic data distributions. **(A–D)** Visualizations of proteomic data reduced to three dimensions, generated by four distinct dimensionality reduction techniques: **(A)** Principal Component Analysis (PCA), **(B)** Partial Least Squares (PLS), **(C)** Uniform Manifold Approximation and Projection (UMAP), and **(D)** t-distributed Stochastic Neighbour Embedding (t-SNE).

#### Full Results of Multiclass Classification Analysis

**Figure S6:**
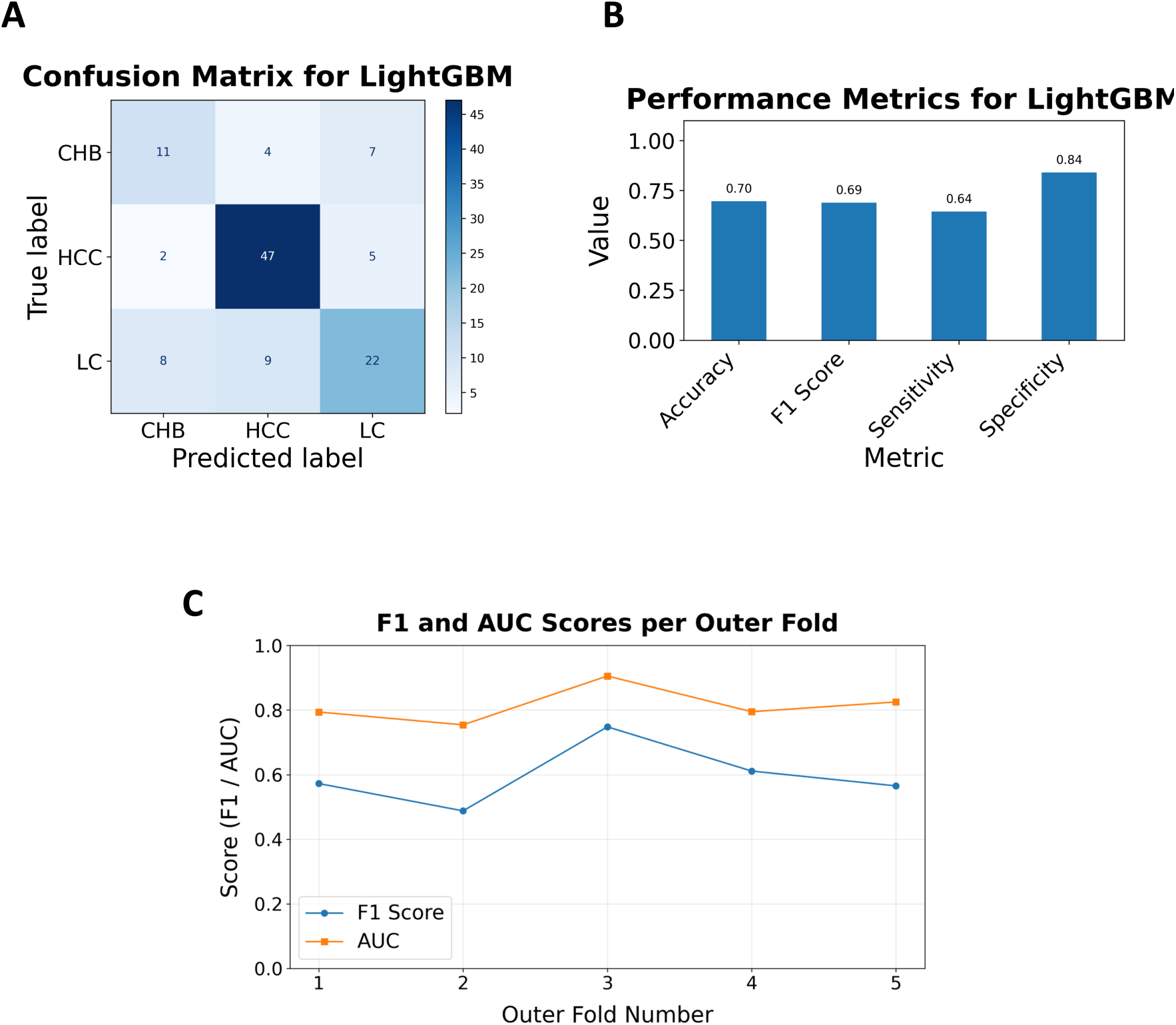
Performance evaluation and model interpretability analyses across liver disease multiclass classification tasks. **(A)** Confusion matrices illustrating per-class prediction accuracy for LightGBM. **(B)** T**he** evaluation metrics, including accuracy, sensitivity, specificity, and weighted F1-score for the classifier**. (C)** Outer cross-validation results showing the distribution of AUC and F1-score across five folds, reflecting model stability.

